# Socioeconomic impacts of elimination of Onchocerciasis in Abu-Hamed focus, northern Sudan: lessons after elimination

**DOI:** 10.1101/858878

**Authors:** Ayman Ahmed, Anas Elbashir, Asgad Adil, Asha A.Alim, Asia Mubarak, Duaa Abdelrahman, Eilaf Mohammed, Noah Saad Mohamed, Arwa Elaagip, Isam M.A. Zarroug, Noma Mounkaila, Hanan Tahir

**Affiliations:** Institute of Endemic Diseases, University of Khartoum, Khartoum, Sudan; Public and Tropical Health Programmes, University of Medical Sciences and Technology, Khartoum, Sudan; Nile University, Khartoum, Sudan; Department of Parasitology and Medical Entomology, Faculty of Medical Laboratory Sciences, University of Khartoum, Khartoum, Sudan; Onchocerciasis Control/Elimination Programme, National Programme for Prevention of Blindness (NPPB), Federal Ministry of Health, Khartoum, Sudan

**Keywords:** Onchocerciasis, black fly, Neglected Tropical Diseases (NTDs), elimination, Mass drug administration (MDA), Socioeconomic, Abu-Hamed, Sudan

## Abstract

**Introduction:** Onchocerciasis is one of the most devastating Neglected Tropical Diseases (NTDs) and it is mostly prevalent in Africa. The disease has important heavy social and economic burdens on the infected populations including low productivity, unemployment, social isolation, and stigma.

**Methodology/Principal Findings:** The socio-economic impacts of the Onchocerciasis elimination in Abu-Hamed, River Nile State, Sudan; were investigated using a well-established questionnaire, 512 participants in ten affected communities were interviewed. Our findings revealed that these communities are recovering from the social and economic burden of the diseases, with 90% of the research participants reported general satisfaction about the elimination of the disease in their community, and about 48.3% of them attended secondary school or university. Only 0.6% reported unemployment, and 25.3% and 24.7% of the participants were workers and farmers respectively. Except about the vector biting and nuisance, the majority of the respondents (90%) had no complain after the elimination of the disease in the area. Also, 90.5% of the participants reported either stable or increase in their work performance during the last twelve months. About 93.8% of the respondents were engaged in normal daily activities and involved in happy events like marriage and giving birth during the last twelve months.

**Conclusions/Significance:** Overall, we conclude that the elimination of Onchocerciasis in Abu-Hamed has several positive impacts on the economy and social life of Abu-Hamed local communities, but this could be maximized by controlling the vector, which is still having a negative impact on the populations. Establishing local developmental projects will help these communities greatly to recover and become more productive.

**Author Summary:** Onchocerciasis, also known as the river blindness, is a disease that caused by a parasitic worm which could infest people eyes or skin causing a blindness or sub-dermal disease. This worm is transmitted to human by the bite of an insect, the black fly. Although the disease is not fatal in most of the patient but it presents a significant economic and social burden on the infected people, their families and communities. This burden is a result of the social stigma associated with the skin form of the disease, and the lack of vision in case of blindness (ocular manifestation). Onchocerciasis was a public health problem in the study site, Abu-Hamed. In 2015, the disease was officially has been declared eliminated from the area. We investigated the socio-economic impacts of this success on the local communities. We have interviewed randomly selected 512 participants to understand their perspective and highlighting their experience regarding the disease elimination. Our research aimed to fill the gap of knowledge between the public health and social science by highlighting the social and economic benefits of health interventions and diseases elimination/eradication. Furthermore, to urge intervention programs to empower the local communities in the planning and implementation of health interventions for a better success.

## Introduction

Onchocerciasis, commonly known as the river blindness disease, is a parasitic disease that is caused by the worm *Onchocerca volvulus*. It is transmitted by the female black flies, mainly *Simulium damnosum sensus lato*, of the genus *Simulium* (1). Onchocerciasis is one of the major neglected tropical diseases (NTDs) and it is mainly prevailing in Africa with confirmed present in 31 African countries, 6 countries in Americas and one Asian country, Yemen, with more than 100 million people at high risk of infection globally (2). The mature worm has long reproductive lifespan up till 9-11 years; period during which it releases numbers of microfilariae (mf) daily in human host body (1,3,4). The Onchocerciasis Control Programs are relying on Community Directed Treatment with Ivermectin (CDTI) distributing the freely donated ivermectin (Mectizan®) to the populations at risk annually or biannually (5,1,3). Unfortunately, the drug only kills the mf but has no lethal effect on the mature worm, however, it might significantly reduce the worm productivity (3). The death of mfs inside the host tissues with repeated infections can cause serious inflammatory immune response and damage to vital organs like skin, eye, and brain (3,6).

Onchocerciasis is the second leading infectious cause of irreversible eye blindness, resulting in 1 million cases (7). Also, it causes a spectrum of serious infections, including skin itching “Onchocercal skin disease” (8), muscles pain and general malaise (9), weight loss, and enlargement of the genital organs (10,11). Also, the severe presentation of the disease is associated with developing epilepsy disease (12,6). In 2015, nearly 117,000 persons were estimated to be affected by the onchocerciasis-associated epilepsy with estimated cost of treatment up to 12.4 million US$ for them (13). The estimated international economic benefit from the averted productivity loss due to Onchocerciasis is 4.4 [3.2–6.9] billion dollars (14). Success stories were achieved in controlling and eliminating different neglected tropical diseases worldwide, and Abu-Hamed is the first focus of Onchocerciasis in Africa to be eliminated and certified by WHO after 3-years post-treatment surveillance (7,15).

The devastating disability associated with NTDs is extremely burdensome, that decreasing the lifestyle of the poorest of the world, their work productivity and socio-economic outputs (7). Onchocerciasis has many social and economic negative impacts in addition to the health effects; it exacerbates productivity, social and sexual life, and consequently leads to poverty that obstacles the development of poor communities (16,7,12,17) with heavy burden of social stigma and isolation which limits their chances in education, labor, wage earning, and normal social life activities, resulting in developing severe psychological disorders (7,9,18). Therefore, controlling and eliminating this disease will develop and enhance the health and lifestyle of the vulnerable populations and directly contributing in achieving the Sustainable Development Goals (SDGs) (7,18,19).

Onchocerciasis found in Sudan in four foci: Northern Sudan (Abu-Hamed focus, Northern State); Eastern Sudan (Galabat sub-focus, Gadaref state), South and Southwest Sudan (Radom focus, South Darfur state) (20), Southern Sudan (Khor Yabous, Blue Nile state) (Sudan National Onchocerciasis Elimination/Control Programme). The Abu-Hamed focus is unique for being the northernmost Onchocerciasis focus in the world (20–23). The control of Onchocerciasis in Abu-Hamed started with an annual community-based treatment with Ivermectin (CDTI) in 1998. In 2006, the Government of Sudan launched an Onchocerciasis elimination policy and switched from annual to semi-annual (six months) CDTI (15). Comprehensive surveys showed low level of transmission in 2007 (24), and complete interruption of the disease transmission in 2011 that was officially declared by the Sudan government in 2012 (15,25). Following WHO guidelines and after a 3-year post-treatment surveillance, the elimination of Abu Hamed focus was officially declared in 2015 (26).

Information on the social and economic impacts of Onchocerciasis elimination are required to assess and evaluate the success and guide the redevelopment of the focus population (26). The present study was designed to investigate the socio-economic impacts of Onchocerciasis on Abu-Hamed populations and assess the social changes in their lives after the elimination of the disease.

## Methods

### Study area

The Abu-Hamed focus is centered around the River Nile in the middle of the Nubian Desert (N 19° 32.49’− 18° 17.29’, E 32° 13.86’ − 33° 55.00’) at an altitude range of 260-920 km. The breeding of black fly’s aquatic stages lasts from October to May when the level of the Nile is relatively stable (27).

### Community survey

The survey was covered 10 randomly-selected communities in Mograte district, the last area that reported the disease in Abu-Hamed focus (Figure 1) during September to October 2015, the questionnaire was well-constructed and screened the demographic, socioeconomic, and environmental information of the selected communities.

**Figure 1.**
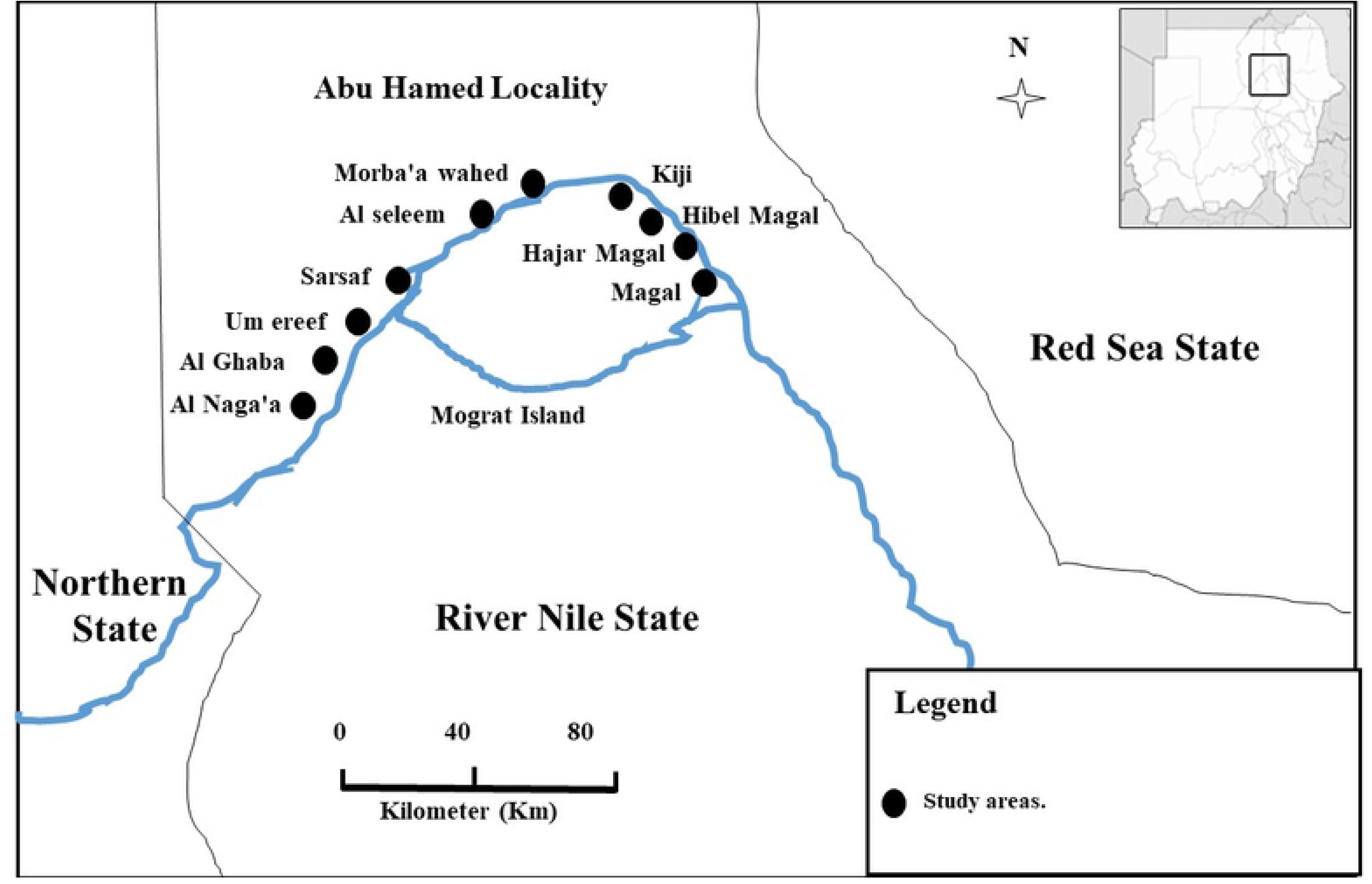
Map display the study sites location in the Sudan.

### Ethical considerations

Ethical approval for the study was obtained from the University of Medical Sciences and Technology and approved by the community leaders, River Nile State Ministry of Health and the Onchocerciasis control/elimination programme. All study participants received full and detailed information regarding the study procedures and objectives using their local language. Each participant agreed verbally and signed an informed consent or their guardians signed to participate as a volunteer in the study.

## Results

This study was implemented in the period of September to October 2015, by the end of the 3-years post-treatment surveillance and the interruption of the disease transmission, and within one year post the declaration of the elimination of the Onchocerciasis in the area. The numbers of research participants from each of the ten selected communities are showed in Table 1.

**Table 1.**
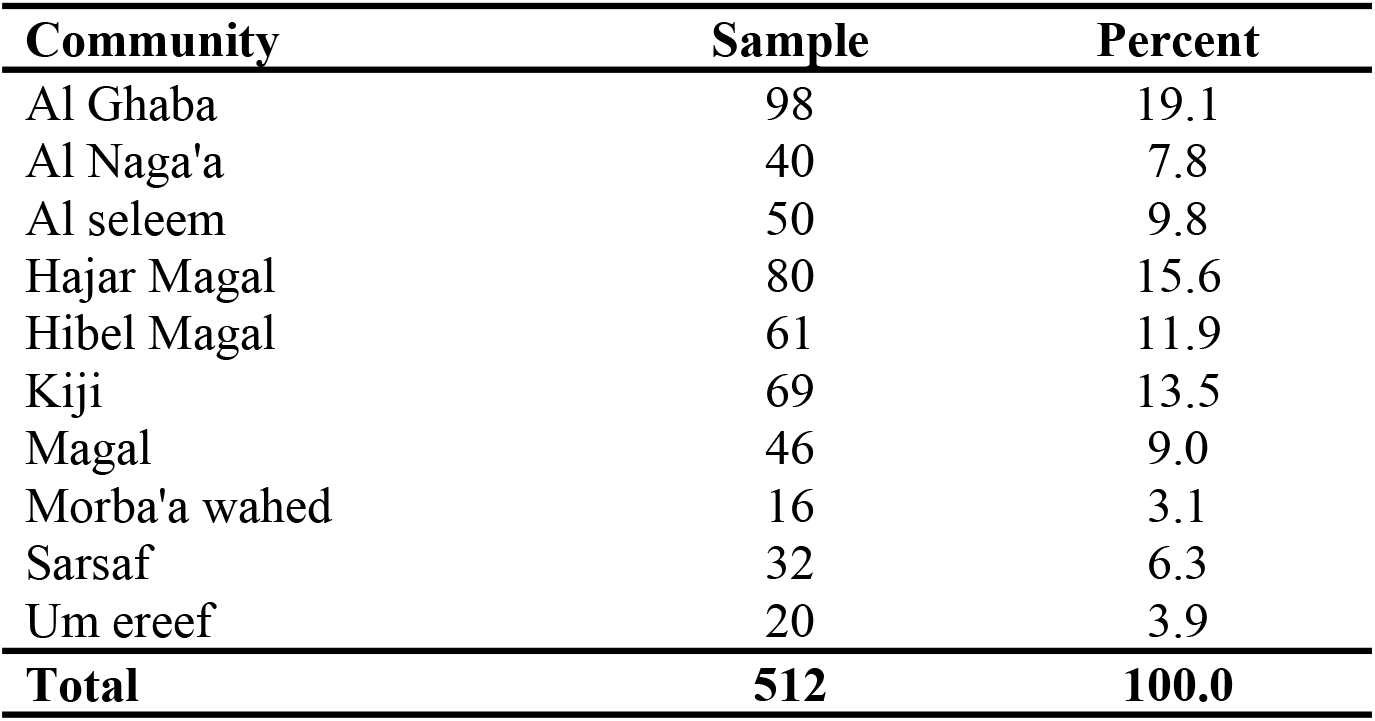
Communities and population selected for the survey in Abu Hamed.

| Community | Sample | Percent |
| --- | --- | --- |
| Al Ghaba | 98 | 19.1 |
| Al Naga'a | 40 | 7.8 |
| Al seleem | 50 | 9.8 |
| Hajar Magal | 80 | 15.6 |
| Hibel Magal | 61 | 11.9 |
| Kiji | 69 | 13.5 |
| Magal | 46 | 9.0 |
| Morba'a wahed | 16 | 3.1 |
| Sarsaf | 32 | 6.3 |
| Um ereef | 20 | 3.9 |
| **Total** | **512** | **100.0** |

### Sociodemographic characteristics of the study population

The study population (n=512) was predominately males with respectively 63% males and 37% females. Fifty-five percent of the participants were male, heads of households, 32.0% were spouses and the remaining 13.0% were daughters/sons, nieces/nephews, brothers/sisters, and other relatives of the head of the households.

The participants (n=512) were 12 to 89 years old with average age of 43 ±16 years. Their duration of stay in their respective communities varied from < 1 year to 89 years with an average year of living in the village of 35 ±19 years. About 89% of the participants were living in their communities more than 10 years, and 82.5% of them did not travel out of their respective communities.

Regarding the education level of the participants (n=509) no answer was obtained from four of them and 12.4% did not attend any form of schooling. Adult schooling was attended by 2.4%, primary school by 36.1%, secondary by 38.7% and university by 9.6% (Table 2).

**Table 2:**
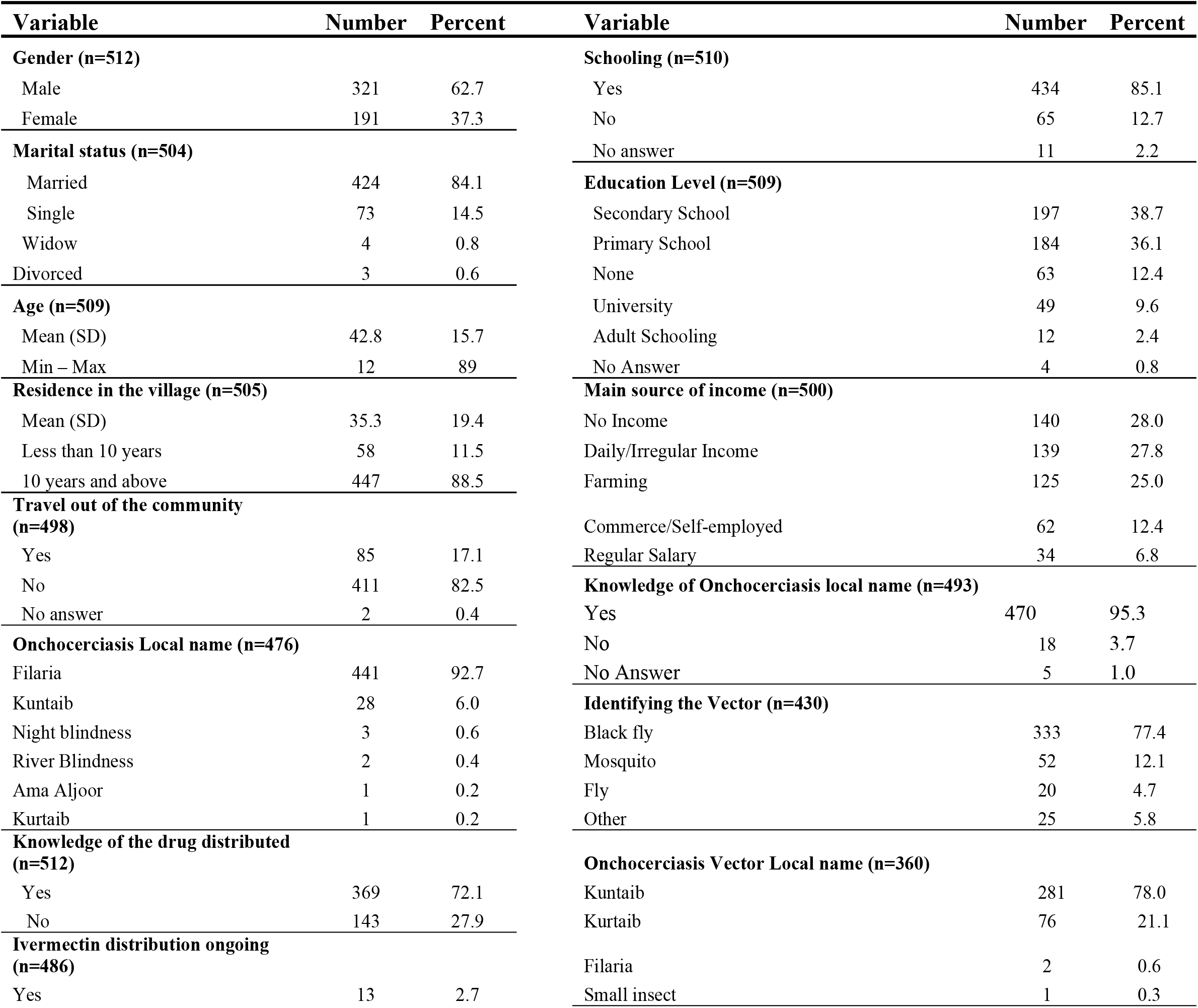
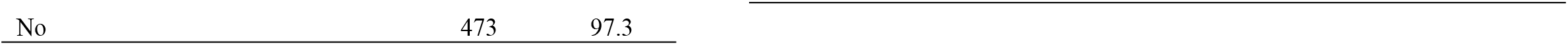
Sociodemographic Characteristics of the study population (n=512)

Unemployment was reported by 0.6% of the participants and 25.9% of the study participants were housewives. The main occupations in Abu-Hamed were laborer (25.3%), farmers (24.7%), and trading/commerce (12.1%). Teachers, drivers and students represented respectively 2.9%, 2.5% and 2.3% respectively. The remaining 4.3% include health professionals (1.9%), security personnel (0.8%), self-employed (0.8%) and a fisher (0.2%). Regarding the main source of income, out of the 500 respondents, 28.0% did not have income. They were either housewives, students or unemployed. Daily or irregular income was recorded for 27.8%. They were mainly labors. Farming was the main source of revenue for 25.0%, commerce/trade or self-employment represent 12.4%. Regular monthly salary was the main source of income for 6.8%; they were either teachers, health professionals, or security forces.

### Health issues and social impact related to onchocerciasis

The participants raised a common compliant in the additional notes section of the questionnaire about the vector biting and nuisance particularly during the winter season when they usually harvesting their farms and crops (Wheat, Okra, and the Egyptian beans) the main source of income in Abu Hamed area.

Most (97.0%, 495/510) of the study population had heard about onchocerciasis, and only 2.8% participants did not hear about the disease. Onchocerciasis is known in Abu-Hamed by several local names which were provided by 95% of the participants (n=493). Onchocerciasis or river blindness is locally named as Ama Aljoor, Night blindness, Kurtaib, Filaria, and Kuntaib. The vector transmitting Onchocerciasis (the *Simulium hamedense*) was identified as black fly by 77.4% of the participants. Surprisingly, the local name of the vector is the same of the disease but not reported in the same proportion (Table 2).

Only 61 participants identified the name of the drug used as Mectizan or Ivermectin^®^. However, 369 participants reported to know the drug and they described it well in terms of presentation and dose administrated by community-distributors. Also, 97.3% of the participants (n=486) reported that ivermectin mass distribution was halted (by the end of project) in their respective communities. The remaining 2.7% who reported continuation of ivermectin distribution had not provided any reason justifying the continuation of the distribution.

Regarding the impact of Onchocerciasis elimination in Abu-Hamed, 90% of the study population (n=512) did not have any complain after the elimination apart from the vector biting and nuisance; persisting itching was reported by 7.2%. Other complains were less than 3.5% and were mainly about the need for developmental project to increase the employability of the locals (Table 3).

**Table 3:** Health issues and social impact parameters.

| Variable | Frequency | Percent | Variable | Frequency | Percent |
| --- | --- | --- | --- | --- | --- |
| **Complain since distribution of the drug was halted (n=512)** |  |  | **Knowledge of people currently complaining about Onchocerciasis symptoms (n=506)** |  |  |
| No complain | 461 | 90.0 | No | 453 | 89.5 |
| Itching | 37 | 7.2 | Yes | 53 | 10.5 |
| Irritation | 13 | 2.5 | **Trend of the work performance of participants (n=377)** |  |  |
| Other | 1 | 0.3 | Stable | 228 | 60.5 |
| **Receiving guests in the twelve past months (n=373)** |  |  | Increased | 113 | 30.0 |
| More often | 167 | 44.8 | Poor | 36 | 9.5 |
| Less often | 128 | 34.3 | **Trend of the revenue of participants (n=371)** |  |  |
| Regularly | 78 | 20.9 | Stable | 226 | 60.9 |
| **Opinion regarding halt of ivermectin distribution (n=482)** |  |  | Decrease | 73 | 19.7 |
| Distribution to continue because of health benefits | 377 | 78.2 | Increase | 72 | 19.4 |
| Programme reached objectives and as halted | 96 | 19.9 | **Receiving guests in the twelve past months (n=373)** |  |  |
| Distribution to continue because of Programme failed | 9 | 1.9 | More often | 167 | 44.8 |
| **Trend in school attendance (n=19)** |  |  | Less often | 128 | 34.3 |
| Increase | 10 | 52.6 | Regularly | 78 | 20.9 |
| Decrease | 7 | 36.8 | **Social events in the last twelve months (n=162)** |  |  |
| Stable | 2 | 10.6 | Birth | 93 | 57.4 |
| **Trend in school performance (n=20)** |  |  | Marriage | 59 | 36.4 |
| Good or better | 12 | 60.0 | Divorce | 9 | 5.6 |
| Fair or worse | 8 | 40.0 | Separate | 1 | 0.6 |

Changes in the performance of children attending school had not been noticed by 91.9% of the study population (n=307), only 19 people (6.2%) noted changes and 1.9% did not know (Table 3). For those who noted changes, school attendance (n=19) increased for 52.6%, for 10.6% the attendance was stable and it decreased for 36.8%. The participants (n=20) rate school performance as excellent (30%) or good (30%); it was fair and poor respectively for 25.0% and 15.0%.

In the last past twelve months, the work performance of the participants (n=377) remained stable for 60.5%, it increased for 30% and was poor for 9.5%. On the other hand, the revenue of the participants (n=371) remained stable for 60.9%; and increased for 19.4%. The remaining 19.7% reported a decrease of their revenue.

Socialization was measured as the frequency of receiving guests and/or getting involved in social events in the last twelve months. Also, 44.8% of the participants (n=373) received guest sometimes, 34.3% less often and 20.9% regularly. In the past twelve months, the social events the most frequent reported by participants (n=162) was birth (57.4%), followed up by marriage (36.4%). Divorce and separation were reported respectively 5.6% and 0.6%.

Participants were requested to share their opinion regarding stopping ivermectin mass distribution (n=482). Additionally, 78.2% thought the distribution should continue because of the health benefits of the drug such as deworming and preventing other skin infections. Only 21.8% was in favor of stopping ivermectin distribution. Out of 105 participants, 91.4% justified the halt of the distribution because of the program objectives were achieved, and failure of the program may justify the decision for the 8.6%.

The 89.5% of participants (n=506) reported that currently at their knowledge no one complains about Onchocerciasis and the remaining 10.5% know people complaining currently about the disease (persistent itching).

In the overall, the social impact of Onchocerciasis elimination in the ten communities surveyed can be estimated with a general public satisfaction reported from most of the study participants (90%).

## Discussion

The Sudanese Onchocerciasis Control/Elimination Program initiated the CDTI in Abu Hamed with annual treatment in 1998 then followed by four years of dual treatment per year from 2007 to 2011, when after assessment and fulfilling the WHO criteria for the disease interruption and stopped the treatment by 2012 (15). We conducted this study at the end of 2015 to assess the socioeconomic impacts of this success and evaluating the community recovery from this disease. The outcome of this evaluation might motivate the stakeholders of the NTDs to invest more in eliminating such devastating diseases.

The fact that Abu Hamed is the most remote focus of Onchocerciasis in the north worldwide with a very limited opportunity of re-establishment of the infection after the elimination (26). This situation provided us with an extraordinary opportunity to study the socioeconomic impacts of the elimination of Onchocerciasis in Abu Hamed focus. Abu Hamed focus of Onchocerciasis was identified for the first time by Morgan in 1958 while the vector was reported in the area five years earlier by Lewis (Haseeb et al., 1962; Morgan, 1958; Lewis, 1953). Clinical studies revealed that this focus was entirely dominated by the dermal form of the disease and no case of the eye form has been reported from this area ever. It might be due to the unique strain of *O. volvulus* circulating in this area (21,29), or the unique sub-species of the vector, *S. hamedense,* which is totally confined to the area (30).

Despite of that, the Ocular manifestation of the disease is the most known form of the disease as it causes blindness and disabling the patients (31). Nevertheless, the dermal form of the disease has more social impacts on patients because it tremendously changes the skin of the patients regarding the color and texture leading to modifying the patient appearance badly (1,32). This in turn results in stigma, family and social isolation, implicating people’s live, and leading to all different type of socioeconomic loses and psychological effects, including unemployment, dropping out of school, less opportunity to marry and establishing a family, and eventually leads mental disturbance (33–37).

Our results showed high rates of recovery at different aspects of the affected population, 11.5% of the study participants (512) were new comers in the area, immigrate within the last 10 years. Though, people usually forced to desert their homes and villages, and leaving to avoid the infection (18,38). Although the percentages of our respondents enrolled in or achieved different level of education seem low, but it is in accordance with the averages of normal rural areas in Sudan (39). Remarkably, only 0.6% of the participants reported unemployment which is much lower than the country overall unemployment rate, which increased to 13% between 2013-2015 (40).

With 90% of the participants (512) had no complain about the disease while the rest complained about itching, social or mental stress either about themselves or someone they knew, this mainly due to previous infection, and they have been referred to proper health and mental care to intervene and help them overcoming their situations. This low percentage might be due to the population engagement with the control program activities which increased their awareness about the disease and reduced the level of stigma (41,42). Although there was no specific question about the vector biting or nuisance in the original questionnaire because the Onchocerciasis control project only followed CDTI strategy and did not deliver any control measure targeting the vector, however a major complaint among the participants made it so important to be highlighted because it affects their work in their farms and fishing or other activities near to the river banks and getting very intense during the harvesting time of their main fruit, the date during the winter (27,43).

In this study, 95% of the working participants, reported improvement or stability in their work performance leading to sustainable income and reducing the risk of poverty (18,34). There was fluctuation in the students’ school attendance. We perceived that as an improvement in their health situation associated with elimination because it correlates with areas most affected by the disease. Suggesting that these students are contributing in the family work to generate income (18).

Majority of our participants (93.8%) reported they were involved in one or more happy event during the last 12 months indicating high rate of social recovery (33,37,44).

In conclusion, the elimination of Onchocerciasis in Abu Hamed area is a success story needs to be shared for inspiration. Apparently, the affected communities are recovering from the burden of the negative social and economic impacts of the disease, but still there is a long way to go for them to overcome some side effects of the disease include achieving the local developmental plans. However, they are still in need for a solution for the vector biting and nuisance which is greatly affecting their productivity.

## Acknowledgments

We thank the local communities for their hospitality and participation in the study. Also, we would like to thank our colleagues at the River Nile State Ministry of Health and the Sudan Onchocerciasis Control/Elimination Programme for their support and help throughout the implementation of the project.

## Compete of interests

Authors declare that they have no competing of interests.

